# Mouse Organ Specific Proteins and Functions

**DOI:** 10.1101/2021.11.01.466788

**Authors:** B. Sun, C. Lorang, S. Qin, Y. Zhang, K. Liu, G. Li, Z. Sun, A. Francke, A. G. Utleg, P. Flores, Z. Hu, K. Wang, R. Moritz, L. Hood

## Abstract

Organ specific proteins (OSPs) possess great medical potentials both in clinics and in biomedical research. Applications of them — such as alanine transaminase, aspartate transaminase, and troponins — in clinics have raised certain concerns of their organ specificity. The dynamics and diversity of protein expression in heterogeneous human population are well known, yet their effects on OSPs are less addressed. Here we use mouse as a model and implemented a scheme of breadth study to examine the pan-organ proteome for potential variations of organ specificity in different genetic backgrounds. Using reasonable resources, we generated pan-organ proteomes of four in-bred mouse strains. The results revealed a large diversity that is more profound among OSPs than the overall proteomes. We defined a robustness score to quantify such variation and derived three sets of OSPs with different stringencies. In the meantime, we found that the enriched biological functions of OSPs are also organ specific that are sensitive and useful to assess the quality of OSPs. We hope our breadth study can open doors to explore the molecular diversity and dynamics of organ specificity at the protein level.

## INTRODUCTION

Multiorgan mammals are evolved in such that each organ has unique morphology and functions. Their molecular underpinnings have been the foundation of diagnosis and treatment of various diseases as well as the understanding of disease etiology. Based on the organ specificity, many molecular assays are developed for monitoring the organ pathological states and disease progression, such as the cardiac trophonin (cTn) test for the heart ^1^, the alanine transaminase (ALT) and aspartate transaminase (AST) tests for the liver ^2^, and the prostate specific membrane antigen (PSMA) for the prostate cancer ^3^. However, these molecular markers are only available for a limited number of organs and diseases, and often times their organ specificity is in question^4^.

Several reasons hinder the research and broad applications of organ specific proteins (OSPs). First is their definition. The organ specificity is in fact challenging to define, particularly when we do not have a good consensus on the exact number and types of organs/tissues in humans. The narrow definition of organ specificity requires the presence in only one but absence in the rest of the organs. If we do not know exactly what the rest organs are, the specificity is relative but not absolute. Second, we still cannot detect all proteins in humans. The current mapping of human proteome is around 90% ^5,6^. Due to the detectability, the absolute specificity of OSPs is elusive. Third and maybe the most pressing challenge is the dynamics and diversity of protein expression, which compounds the first two challenges. The dynamics ─ we define here ─ is the change of protein expression across time of the same individual; whereas the diversity here is the change across different genetic backgrounds in a population. A snapshot of a proteome cannot represent the whole. For example, many large scale and deep proteome characterizations on humans ^7,8^ as well as on animal models such as mice ^9,10^ reported a single ensemble proteome. When studying organ specificity using such an ensemble approach, the dynamics and diversity of the proteome are averaged. Yet, pathological conditions, for example, are accompanied by changes of protein expression that can interfere protein organ/tissue specificity and their usage as disease diagnoses. In The Cancer Genome Atlas project (TCGA) ^11^, numerous novel genes were detected in ectopic organs, such as the cardiac troponin I in lung cancer tissues ^12^.

To address the first two challenges, more tissue and organ types have been examined and more sensitive methods were employed for higher coverage and deeper penetration of the proteome. But the mounting cost, time, and effort prevented such depth studies to include broad conditions and genetic backgrounds. In the other front, genetic diversity has been recently explored at large scale through association studies in both transcriptomes and proteomes ^13-15^, i.e. eQTLs and pQTLs. Even though these studies tend to focus on a single tissue type or simple organisms, a large variation has been reported. Community efforts are developed to address the diversity at multi-tissue levels such as the GTEx ^16^, ENCODE ^17,18^, TCGA ^11^, and the Human Proteome Project (HUPO) ^6^ and Human Proteome Atlas ^8^. Due to the immense data, analyses often require highly experienced bioinformaticians with sufficient details on both the study and the data for meaningful results. At proteome level, the diversity and dynamics of the organ proteins are well known but are less systematically evaluated than those of transcriptomes.

As a result, for clinically used organ proteins such as ALT and AST, none of them are absolutely specific to their target organs ^4^. Both ALT and AST have leakage expression in a number of other organs, such as the kidney. However, clinicians can use complementary assays to provide accurate diagnosis ^19^, once the differential expression of ALT and AST between the hepatic and extrahepatic organs are known. Therefore, even though ALT and AST are not *bona fide* liver-specific proteins, the knowledge of their blood concentration in normal and pathological conditions still allowed them to be the popular clinical tests for liver functions. In the early pandemic of Covid 19, a lack of understanding on the precise expression of ACE2, the virus main target receptor, had challenged health professionals to comprehend the mosaic pathological phenotypes observed in patients. By pooling known information together, it is evidenced that ACE2 is ubiquitously expressed in all tissues by transcriptomics, but localized only to a number of specific cell types and body parts by direct protein measurements^20^. How would diseases, such as pre-existing conditions, shift such expression pattern, remains poorly addressed. Therefore, to understand and to treat a novel disease will take us a long time if unprepared. Furthermore, we probably far less equipped to diagnosis and treat other diseases that are less studied than Covid.

We are urged to ask how diverse protein expression can be, and to what extent the information on protein localization or tissue specificity can be clinically useful tools for diagnosis and treatment of diseases. A quantitative evaluation will be more informative than qualitative understanding, especially in clinical setting when both clinicians and patients demand to know the effectiveness of a particular diagnosis or treatment. To our knowledge, no quantitative evaluation has been made to multiorgan diversity of protein expression across different genetic backgrounds, even though for particular protein biomarkers, the sensitivity and specificity are frequently evaluated in large patient cohorts. To fill the gap, we here use mouse as a model and studied the proteome of 13 major organs from the commonly used BL6 strain and verified the selected organ proteomes in three other in-bred mouse strains, i.e. A/J, SJL, and NOD. We named our study “breadth study” to differ from existing “depth studies”. Our results suggested that some but not all OSPs can vary substantially in different genetic backgrounds. We quantified such changes by robustness scores and diversity scores. Our framework can be easily scaled up to include more conditions, and the analyses can remain simple and straightforward.

## MATERIALS AND METHODS

Mouse strains of C57BL/6, A/J, SJL, NOD were purchased from Jackson Lab. Protease Inhibitor Cocktail was from Sigma. Tris(2-carboxyethyl)phosphine (TCEP) and BCA protein assay kit were from Pierce and sequence-grade trypsin was from Promega. Rapigest and Sep-Pak C18 columns were from Waters and C18 Zip tips were from Millipore. All other chemicals were purchased from Fisher Scientific.

### Harvest of animal organs

The animal studies have been approved by Institutional Animal Care & Usage Committee (IACUC) (10-00 series) of Institute for Systems Biology (Seattle, WA) with an assurance from the Office of Laboratory Animal Welfare (OLAW Assurance no. A4355-01) and been accredited by the Association for Assessment and Accreditation of Laboratory Animal Care (AAALAC Accreditation no. 001363). All methods were carried out in accordance with relevant guidelines and regulations. Eight-to nine-week-old mice were fasted for 24 hours prior to the organ collection. Animals were euthanized and dissected immediately for the targeted organs and tissues, including the liver, lung, kidney, heart, brain, small intestine, eye, pancreas, spleen, bone, skeletal muscle, fat, and the testis. Sections of the excised organs were stored in 10% formalin for H&E staining in the Department of Pathology, University of Washington. Details are in the Supplementary material. The rest organs were snap frozen in liquid nitrogen and stored in -80 °C for proteomic analysis. The study was in compliance with the ARRIVE guidelines.

### Proteomic sample processing

Frozen organs were pulverized in ceramic crucible chilled with liquid nitrogen and proteins were extracted with 2-5% SDS in PBS with 1:100 diluted Protease Inhibitor Cocktail by Precellys24 (Bertin, France) at 4 °C. The extracted protein solutions were digested by trypsin based on the filter aided sample preparation (FASP) method ^21^. The digested sample were desalted on a Sep-Pak C18 column and dried in a SpeedVac^®^ (Thermo Savant, Holbrook, NY, USA) concentrator. Supplementary material describes the details of these steps.

### Analysis of peptides by mass spectrometry

The cleaned and dried peptides were reconstituted with MS loading buffer (0.1% formic acid and 1% acetonitrile), and quantified by nanodrop for peptide concentration. Two micrograms of sample at 1ug/ul concentration was loaded onto nanoLC for ESI-MS/MS analysis based on published procedures ^22,23^. Details are in the supplementary material.

### MS/MS database search, inference and quantitation of sample proteins

MS native data files in different proprietary formats were converted to mzXML or mzML using ProteoWizard msconvert ^24,25^ and the spectra having fewer than 6 ions with intensity less than 100 were discarded ^26^. MS/MS spectra were searched with Sequest ^27^ against a Uniprot Complete Proteome database, comprising 51551 mouse proteins, common contaminants listed in the common repository of adventitious proteins (cRAP) and a sequence-shuffled decoy counterpart were appended to the database. Detailed search parameters can be found in the supplementary material. The search results were processed with the Trans-Proteomic Pipeline (TPP, version 4.7) ^28^ including PeptideProphet ^29^, iProphet ^30^ and ProteinProphet ^31^, in which technical and biological replicates of the same tissue were combined. A false discovery rate of less than 0.05 was used to filter the results at protein level. We then estimated protein expression by spectra counting ^32^. Proteins were normalized within each tissue by the maximum spectra count ^33^.

### Analysis of organ specific proteins

We used detection breadth (DB) to describe the distribution of each protein in the sampled organs. Based on DB, we defined our organ specific proteins (OSPs) as DB = 1, i.e. proteins detected only in one organ without specific descriptions. For the rest proteins, we defined common (DB = all) and shared (all > DB **≥** 2) proteins. When verifying how each protein behaved across different mouse stains, we defined the Robustness score (the R score), for which the detection in a distinct genetic background was scored 1. Therefore, the maximum R score in our study is 4. Because we frequently compared proteins across multiple strains or conditions, we also defined the Diversity score (D score) that was normalized between 0 ∼ 1 for the degree of difference between the compared individual to the whole. The D score is computed by

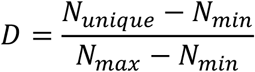

the N_unique_ is the number of unique entries compared across datasets, N_min_ is the minimum possible number of unique entries, and N_max_ is the total number of combined entries before removing the redundancy. The maximum value of the D score is “1”, which means all entries are unique to each other and no overlap exists. A value of less than 1 suggests that certain entries are shared by two or more datasets, i.e. a decreased diversity. A value of “0” means identical entries are among all datasets, therefore no diversity.

### Functional enrichment analysis

We used DAVID Bioinformatics Resources ^34^ to analyze the enriched biological functions in our datasets based on the Gene Ontology Biological Processes (GO_BP). Using the default EASE score (a modified Fisher exact p-value that is more conservative) higher than 0.05, we obtained the enriched GO_BP terms against the total mouse genome background. The definition of organ specific functions was by the presence in only one type of organs/tissues.

## RESULTS AND DISCUSSION

### Overall results

In our study, we applied shotgun proteomics and characterized relatively abundant proteins from 13 major organs in C57BL/6J (BL6) as denoted in **Fig**. 1A, i.e. the brain, spleen, eye, bone, fat, lung, heart, kidney, liver, intestines, pancreas, testis, and the muscle. We then examined the selected organ proteomes from three other in-bred mouse strains including A/J, NOD/ShiLtJ (NOD), and SJL/J (SJL). All the results were deposited in PeptideAtlas (http://www.peptideatlas.org).

**Fig. 1.**
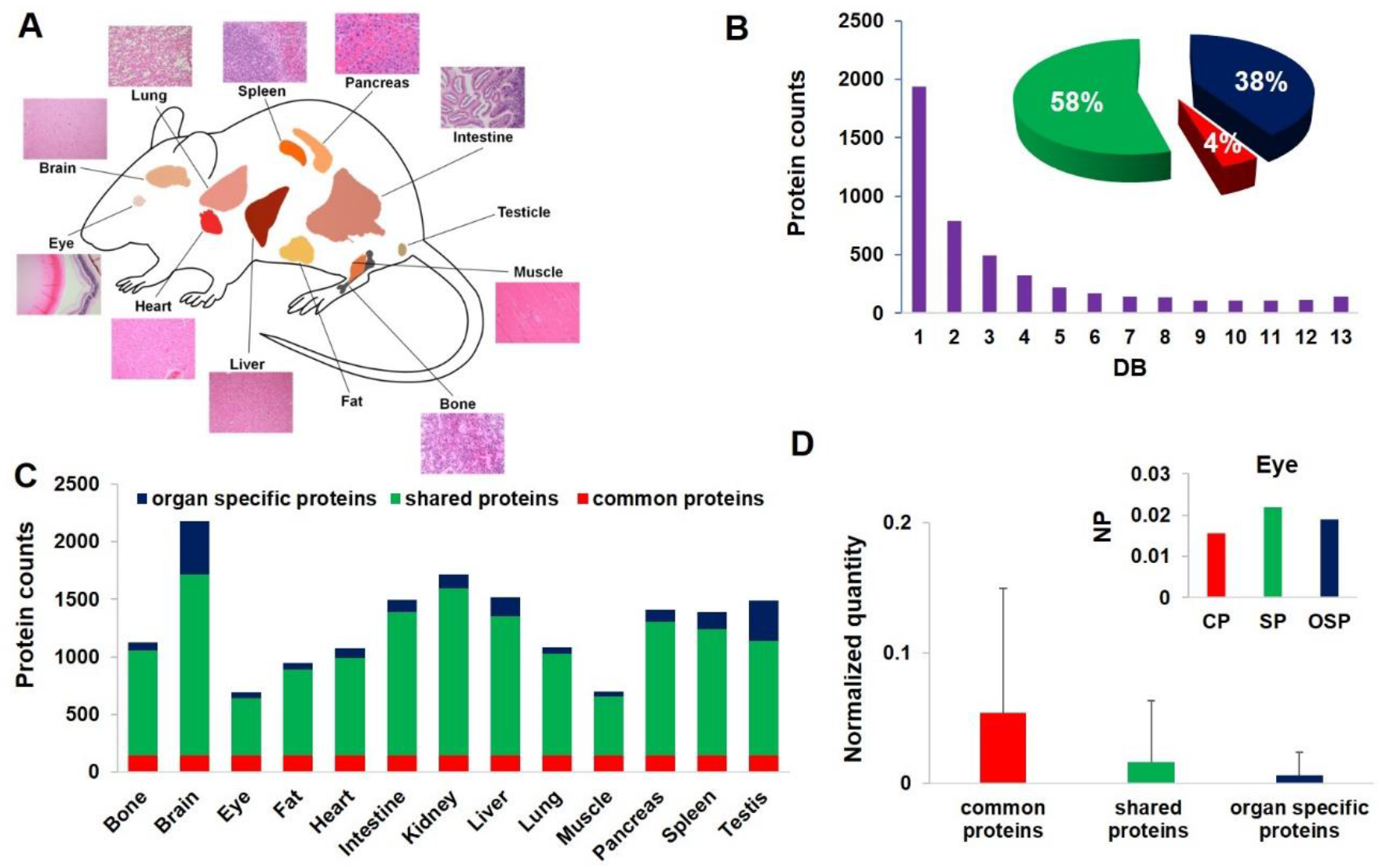
Overall results of mouse pan organ proteome. **A)** Illustrations and histology of the organs and tissues have been analyzed. **B)** The distribution of detected proteins as the function of detection breadth, i.e. the number of organs/tissues, in which a protein was detected. The insert is the percentage of common proteins (DB=13), shared proteins (DB=2∼12), and organ specific proteins (DB=1) in the total proteome. **C)** The number of proteins detected per organ/tissue. **D)** The averaged and normalized detection quantity of the three groups of proteins, error bar is the standard deviation. The insert is the protein detection quantity in the eye, in which NP is Normalized Quantity, CP is Common Proteins, SP is Shared Proteins, OSP is Organ Specific Proteins.

To be practical, we reduced the analysis depth by avoiding the additional off-line fractionations, and only used a single LC-MS/MS injection for each organ. Such analysis only detects relatively abundant proteins, but allowed us to achieve desired organ and strain coverage with reasonable instrumental time. Comparing to the depth studies, we only identified 4787 protein groups from 163 MS runs (Supplementary Table S1). This result is far fewer than the 7349 proteins detected by Geiger et al. from 28 tissue types of BL6 ^9^ with deep fractionation and a total of 336 MS runs. However, we were able to extend the proteome coverage from BL6 to three more genetic backgrounds. Such expansion is crucial for us to evaluate the variability of organ specific proteins (OSPs).

### Protein distribution across tissues

We used the detection breadth (DB), i.e. number of organs/tissues from which a particular protein was detected, to describe the detection distribution. **Fig**. 1B summarizes the result, in which the number of identified proteins displays a concave-shape as the function of DB. A trend is consistent with reported studies on human and mouse proteomes and transcriptomes ^7,9^,35. Using DB value, we defined three types of proteins, i.e. organ specific proteins (OSPs) (DB=1), common proteins (DB=13), and shared proteins (DB=2∼12). The percentage of proteins in each category to the total proteome is shown in the **Fig**. 1B insert. When the total identification was broken down to individual organs shown as a bar chart in **Fig**. 1C, the brain and the eye have the highest and lowest number of the detected proteins, respectively, which have been consistent with previous reports ^36^. We estimated protein expression by normalized spectra counting ^32^. Common proteins have on average a higher expression than shared proteins and OSPs as shown in **Fig**. 1D. When breaking it down to individual organs, the trend is the same except for the eye as shown in the **Fig**. 1D insert. For the eye, many OSPs such as crystallin A are dominant and show higher expression than abundant common proteins, such as alpha collagen, beta actin, and alpha enolase, suggesting OSPs are not necessarily always in low abundance. It is therefore important to catalog OSPs for their individual characteristics.

### Organ specificity of proteins

As we introduced that organ specificity is relative to the detected proteins, and the search of the absolute organ specificity is currently impractical. Yet, with consistent analysis depth and uniform sample processing scheme, we can define relative organ specificity or conditional specificity. Because of our relatively shallow penetration in the proteome comparing to the depth study, our detection of leakage expression is less severe. As a result, more OSPs are defined in our dataset. For example, 1602 OSPs were derived in BL6 that occupied 40% of the total identified proteins; whereas only 43 proteins, i.e. 1.1% of the total identification, were commonly detected in all organs examined. This result contrasts those of depth studies. Specifically, in the recently mapped human proteome, more than 2350 proteins (∼13.6%) ^7^ were detected as common; and in the BL6 mouse proteome, 1972 proteins (27%) ^9^ were detected ubiquitously and only 650 proteins (8.8%) were from one tissue. Clearly higher penetration in the proteome expanded the pool of common proteins, even though many of them are dominant in only one or a subset of tissues. By sampling relatively abundant proteins, we effectively avoided the detection of leakage proteins that are present in low abundance ubiquitously, which helped to increase the OSP population. However, the shallow penetration will miss many critical low-abundance proteins that are also organ specific, such as surface proteins and transcription factors.

Next, we studied how these OSPs change in different genetic backgrounds. For this, we extended the analysis in BL6 to the additional three strains, i.e. A/J, SJL, and NOD. The total trend in terms of the percentage of OSPs and common proteins to the respective total proteome of each strain is very similar. When we compared the pan-organ proteomes among the four strains, the Diversity score is 0.25. When we compared the OSPs, however, the D score is 0.63, which is much higher than the overall proteomes. This observation suggests the OSPs are more sensitive to the diversity of the genetic backgrounds than the total proteomes.

Taking advantages of the multigenetic backgrounds, we filtered our OSPs for their robustness. First, we excluded any OSPs that were designated to different organs/tissues among the four strains, which were more frequent in organs that are closely associated, such as the heart and the muscle. Next, we scored the rest of OSPs with a R value, i.e. the frequency of detecting the same organ specificity among the four strains. The maximum R value is 4, and we filtered the OSPs with R **≥** 2. A criterion indicates an OSP needs to be verified in at least two out of four mouse strains. A total of 655 proteins were passed the filtering, and we named them Verified OSPs (vOSPs) as listed in Supplementary Table S2, including 467 proteins with R = 2, 151 proteins with R = 3, and 37 proteins with R = 4.

To compare the biological significance of the vOSPs from our breadth study to depth studies, we analyzed the functions enriched by the vOSPs. **Fig**. 2A and Supplementary Table S3 summarize the top 10 most enriched biological processes obtained by DAVID Bioinformatics Resources ^34^. It is clear that the enriched biological processes can well represent the organ functions, an observation which is consistent with those of the depth studies ^8,9^. We further examined the organ specificity of the enriched biological processes. The D score for the enriched biological processes is 0.84, suggesting high organ specificity. Some of these characteristic functions are listed here. In the eye vOSPs, “eye development”, “visual perception” and “lens fiber cell differentiation” are top hits; those for brain vOSPs include “nervous system development”, “synaptic signaling” and “neurogenesis”; those for bone vOSPs include “skeletal system development”, “bone development”, “cartilage development”, “extracellular matrix organization”, and “ossification”; those for the lung comprise “cell adhesion”, “regulation of defense response”, “regulation of response to external stimulus”, and “blood vessel development”; those for the heart mostly share with those from the muscle including “muscle cell development”, “striated muscle development” and “muscle system processing”; those for the kidney include “ion transport”, “transmembrane transport”. Several metabolic processes enriched in the vOSPs of both the kidney and the liver, such as “organic acid metabolic process”, “carboxylic acid metabolic process”, and “oxoacid metabolic process”. For liver vOSPs, the uniquely enriched biological processes are “lipid metabolic process”, “monocarboxylic acid metabolic process”, and “fatty acid metabolic process”. The only organ we did not verify is testis as it has relatively large number of unique OSPs as previously reported ^35^, which can be reflected by their enriched GO terms: “sexual reproduction” and “gamete generation”. We did, however, use the testis proteome to eliminate OSPs of other organs as it expresses vast types of different proteins.

**Fig. 2.**
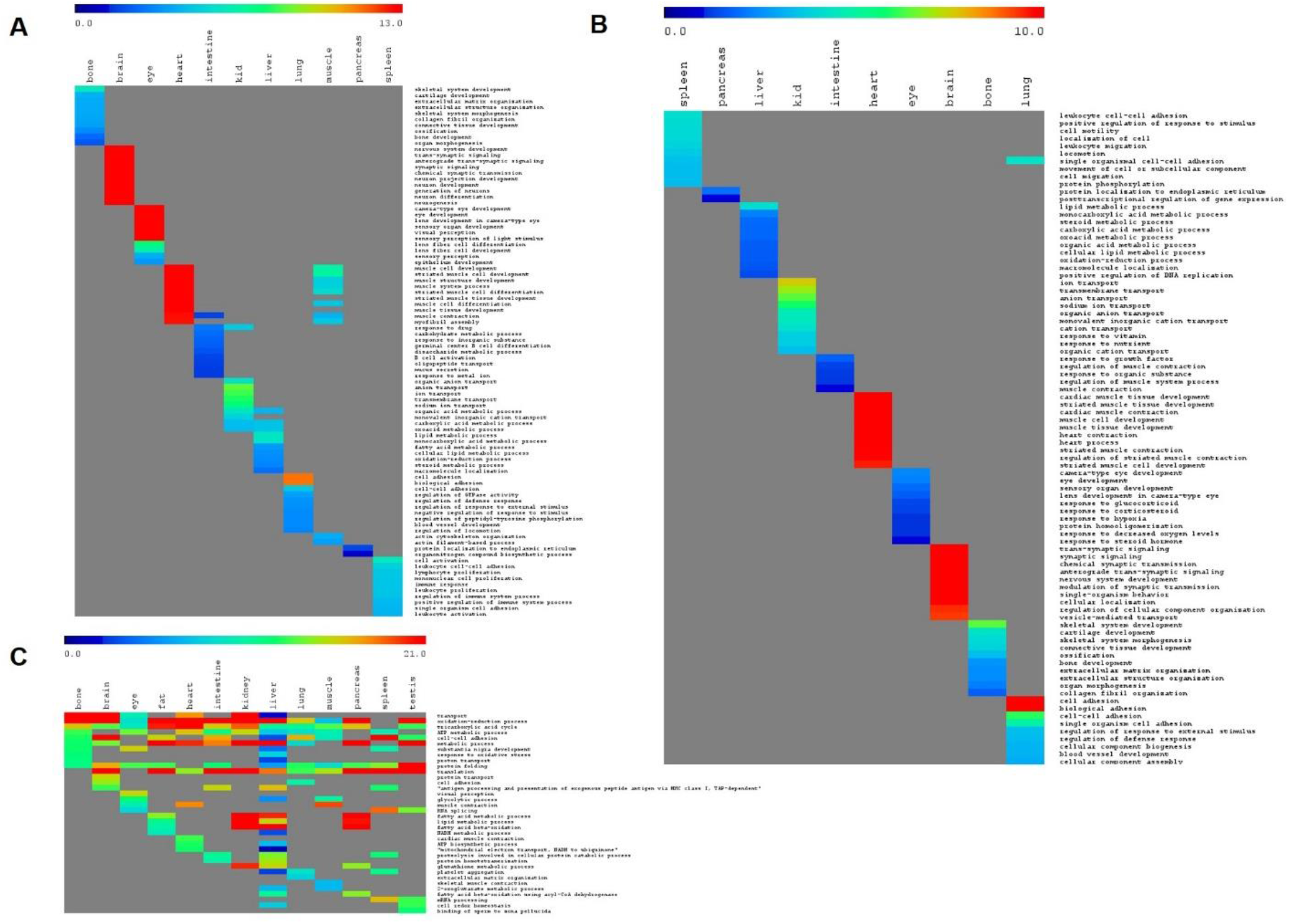
Gene Ontology enrichment analysis of biological processes in the obtained protein lists. **A)** Top10 enriched biological processes of filtered and partially verified Organ Specific Proteins (vOSPs) of each organ against four genetic backgrounds. **B)** Top10 enriched biological processes of complete OSPs of each organ across four genetic backgrounds. **C)** Top10 enriched biological processes of each organ proteome of BL6.

We further examined those vOSPs with R score < 4. These are vOSPs that were verified in more or equal than two strains but were absent from one or two of the other strains. The reason this occurred is because some proteins were detected in more than one organ in a particular strain, therefore they were absent from the OSP lists of that strain. As a result, these vOSPs only maintain organ specificity to partial genetic backgrounds but not to all.

To search the OSPs that are ubiquitously organ specific, we further filtered our vOSPs to remove these partial verified entries. In the end, only 328 OSPs remained (Supplementary Table S4), and we named them as complete OSPs (cOSPs). Among them, the category of R = 4 remains the same, whereas the number of proteins with R = 3 reduced to 99, and the proteins of R = 2 reduced to 192. A similar functional enrichment analysis showed the highest organ specificity among all the datasets we examined, with a D score of 0.99. The results are summarized in **Fig**. 2B and Supplementary Table S5, in which only one term overlapped between the spleen and the lung, whereas all the other terms are unique to their represented organs. Such filtering can effectively remove overlaps between closely associated organs such as the heart and the muscle as shown in **Fig**. 2A. However, the high stringency renders some organs do not resolve any cOSPs or only a few. For example, the muscle has only 3 proteins in cOSPs, including Junctophilin-1 (Jph1), a member of junctional membrane complexes that is expressed specifically in the triad of skeletal muscle in young mice linking the transverse tubule with sarcoplasmic reticulum membrane ^37,38^. Ectopic expression of Jph1 in the mouse heart caused abnormality in junctional membrane structure ^39^, suggesting its muscle specificity.

### Effects of detection depth and breadth to the OSPs

To verify whether the observed tissue specificity in OSP functions was also represented in the total proteome of each organ, we run similar enrichment analysis to pan-organ proteomes. The top 10 enriched biological processes from each organ of BL6 are summarized in **Fig**. 2C. The inclusion of common and shared proteins largely shifted the pattern of enriched GO terms away from those of OSPs (**Figs**. 2A & 2B). The similarity in biological processes across organs is greatly increased whereas the organ specificity has been diminished. The D score of the obtained biological processes is only 0.22, whereas the D score of those OSPs is 0.88, i.e. a 3-fold increase of organ specificity among the biological processes in OSPs as compared with the pan-organ proteomes. Several GO terms, such as “metabolic process”, “ATP metabolic process”, “translation”, and “protein folding” are broadly shared in all organs.

As previously mentioned, comparing to depth studies, our breadth study is not effective in identifying trace proteins in all organs. As a result, our data has more OSPs and less common proteins. Such shallow penetration of pan-organ proteome can introduce less authentic OSPs, yet our results demonstrated that filtering across strains can effectively eliminate some of these false positives. The above addressed GO enrichment analysis showed that the filtered OSPs can well reflect on organ specific functions. Further comparisons of the enrichment analysis among total proteomes, vOSPs, and cOSPs suggest that the functional enrichment results are sensitive. The inclusion of common or shared proteins can skew the results, and our D score can capture such skewing quantitatively.

Another drawback of shallow penetration is to miss some of the extremely low abundance yet organ specific proteins. To evaluate the potential outcome of such effect, we examined our top100 proteins in each organ. The reason to analyze such proteome is because previous depth studies have demonstrated that this proteome even though abundant can effectively delineate organ and sample type differences ^8,9^, suggesting such proteome has sufficient organ specific features that can be used to establish organ or sample identity. On one hand, because these proteomes are rich of abundant proteins, studying such proteome will reflect on the potential issues linked to shallow sampling. On the other hand, because many proteins in this list are widely detected in multiple organs, it can also reflect on the issues in depth studies in which common proteins prevail.

We analyzed the top100 proteins from each organ for OSPs similarly as how we analyzed the pan-organ proteomes. Considering the top100 proteins in each organ are more commonly detected and shared in multiple organs, we filtered the results for conditional OSPs that were dominantly detected in one and less detected in the rest organs. This criterion is relaxed than how we derived the overall OSPs, in which the proteins were filtered by BD = 1. Specifically, we created two indices for analyses. One is to index the frequency a protein was present in the top100 lists ─ “Index-Top100”. The other is to index the frequency a protein was present in the overall organ proteomes ─ “Index-Overall”. **Fig**. 3 shows both indices as a function of Distribution Breadth (DB). In the figure, the Index-Top100 recapitulates the overall protein distribution in **Fig**. 1B; whereas the Index-Overall shows a much even distribution than Index-Top100. The fatter overall distribution suggests these proteins are widely shared across organs; yet, most of them are not the top100 in most organs as suggested by their value in Index-Top100. This observation suggests the possibility to derive OSPs from Top100 proteomes as they still carry organ difference regardless of their relatively high abundance.

**Fig. 3.**
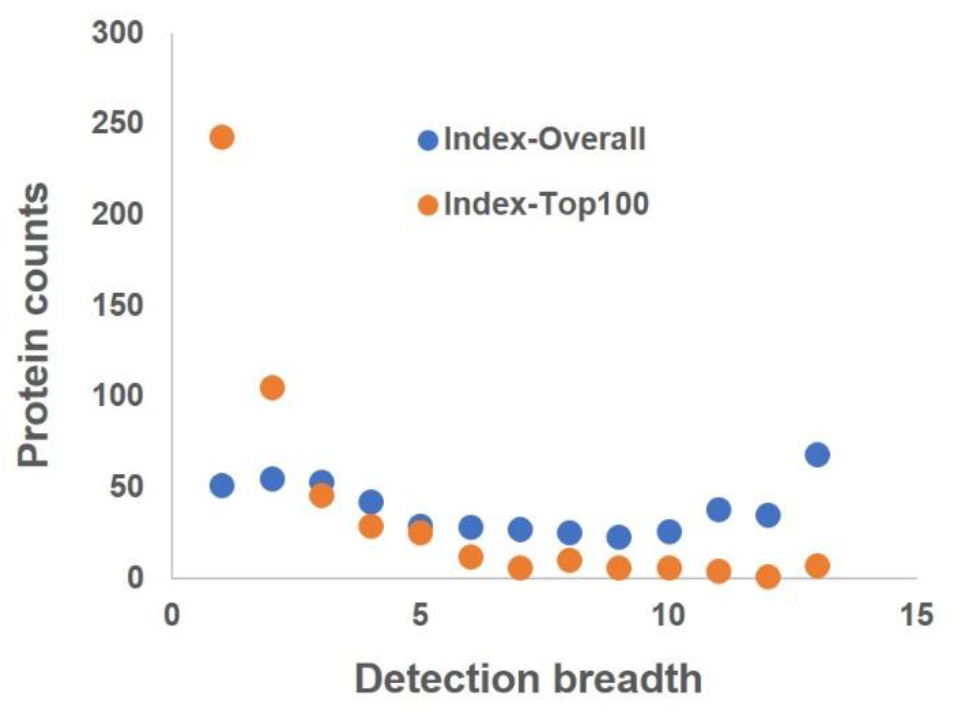
Protein distribution as a function of detection breadth in the organ proteomes, in which the Index-Overall represents the frequency a protein was detected in the overall pan-organ proteome, whereas the Index-Top100 represents the frequency a protein was in the pan-organ top100 proteome.

To derive OSPs from Top100 proteome, we used Index-Top100 = 1 as the criterion to filter the list. To ensure these proteins are dominantly detected in only one organ, we further filtered the results based on the detection quantity difference between the dominant organ and the rest, which we named it as “Dominance Fold”. Using Dominance Fold **≥** 4 as cutoff, we obtained the conditional OSPs from top100 proteomes of all four strains. Next, we examined the protein variation across the strains. Using the R scoring system, we filtered the data similarly by R **≥** 2, and manually verified the obtained proteins for the single-organ dominancy and for the same-organ consistency across genetic backgrounds. In the end, a total of 79 conditional OSPs from the top100 lists were obtained (Supplementary Table S6), and in which 26 proteins are R = 2, 32 proteins are R = 3, and 21 proteins are R = 4.

The conditional OSPs obtained from top100 proteins are compared with vOSPs and cOSPs obtained from the overall proteomes. Clear distinctions exist. First, conditional OSPs are much more abundant in measured quantity than the vOSPs and cOSPs (∼ 2-3 folds higher). Second, the number of proteins from conditional OSPs are fewer than those of vOSPs (> 7-fold reduction) and cOSPs (> 3-fold reduction). Third, the enriched biological processes are not as distinct as those of vOSPs and cOSPs. Fourth, owing to the more common detection among organs, the derivation of conditional OSPs is more challenging and requires more manual evaluation as described above. Out of the 79 conditional OSPs from the Top100 proteome, 43 of them are also part of the vOSPs, but only 15 of them are cOSPs. A result suggests, to obtain adequate number of complete OSPs, certain sampling depth is necessary; yet when the sampling depth becomes too high, the method identifies more leakage expression which can complicate statistical analysis and exponentially increase the cost and resources spent by the experiments.

Besides distinctions, some similarities were also revealed when we compared the three filtered lists of OSPs, i.e. vOSPs, cOSPs, and conditional OSPs from top 100 proteins, against four genetic backgrounds. First, all three filtered OSPs are more abundant in the average detection quantity than the unfiltered lists. For vOSPs and cOSPs, such increase ranged between 3% to 58% in four strains; for conditional OSPs the increase ranged between 25% to 127%. A result suggests random and erroneous detections with low detection quantity can be effectively removed. Second, all filtered OSPs exhibit strain diversity even though much less than the unfiltered counterparts. The D scores for vOSPs, cOSPs, and conditional OSPs of top 100 are 0.24, 0.19 and 0.12, respectively, whereas the D score of the unfiltered OSPs (DB = 1) of the total organ proteomes is 0.63 (63%) and 0.48 (48%) for the unfiltered Top100 (Index of Top100 = 1). This result matches a recent study of liver proteome across a large population of out-bred mice, in which 50% protein diversity was observed among 192 mice ^13^. Our observation suggests that the protein diversity caused by genetic backgrounds is substantial and ubiquitous even for the abundant proteins, and cautions should be kept when use proteins for organ specific applications.

### Organ specific functions

As described above that our GO enrichment analysis demonstrated that the functions derived from OSPs are also organ specific. To further investigate the relationship between the enriched biological functions and the corresponding protein lists, we used BL6 as an example and examined its top10 enriched biological processes from top100 proteins of each organ. **Fig**. 4A and Supplementary Table S7 summarized the results. To compare, the top10 enriched biological processes in OSPs of BL6 are also plotted in **Fig**. 4B. It is clear that the top100 proteins resemble the pan-organ proteomes for the enriched biological processes as shown in **Fig**. 2C, in which shared GO terms across tissue types are common, and dissimilar to those of OSPs in **Figs**. 2A, 2B and 4B. The trend can be well reflected by the D scores across different organs as well. The D score of the enriched biological processes for the Top100 of BL6 is 0.34; those for the total proteome and OSPs of BL6 are 0.22 and 0.88, respectively. The higher D score in the Top100 proteomes than that of total proteomes suggests that organ specific functions are also represented by abundant proteins besides low abundance ones.

**Fig. 4.**
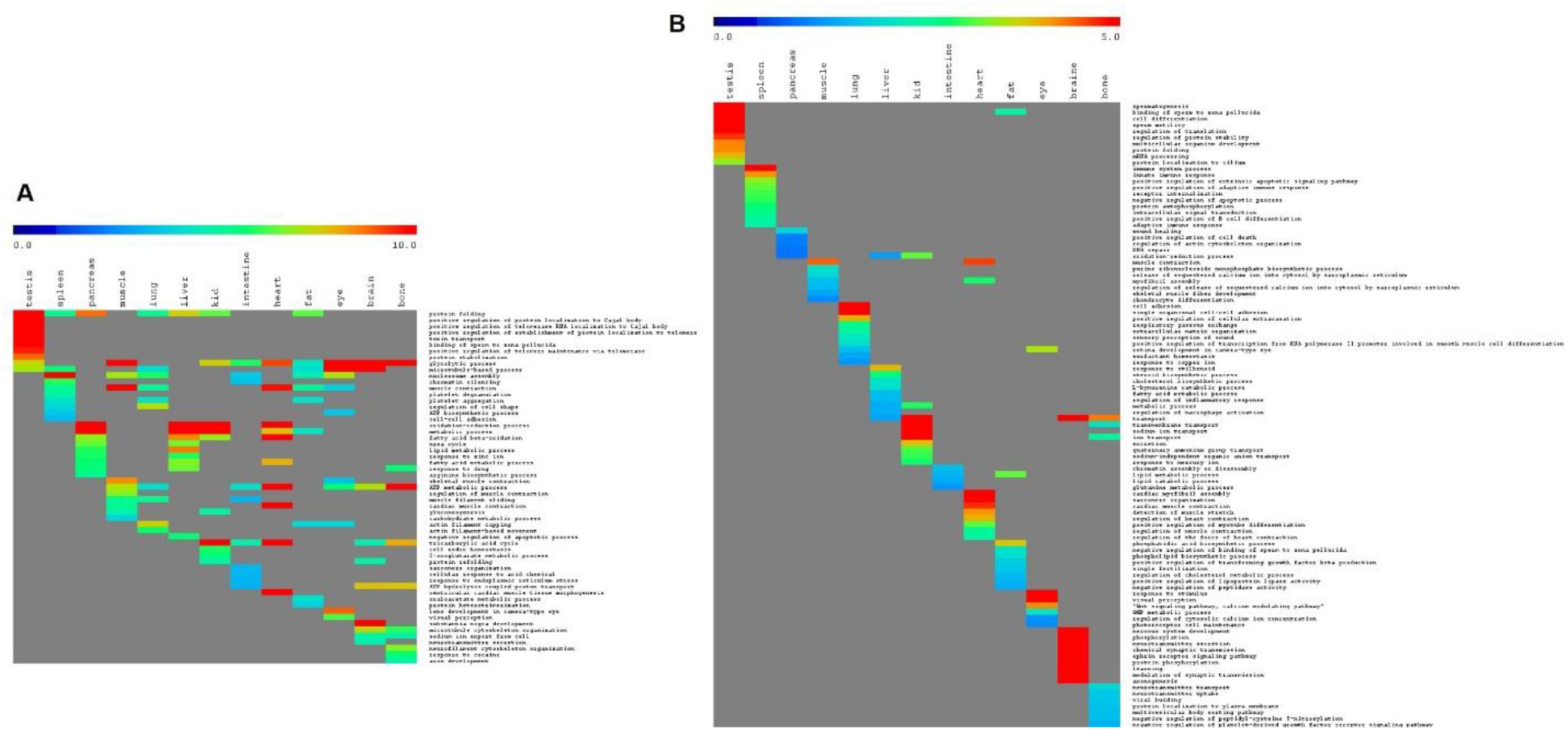
Gene Ontology enrichment analysis of biological processes in the pan-organ proteome of BL6. **A)** Enrichment analysis of the top100 pan-organ proteome. **B)** Enrichment analysis of the Organ Specific Proteins.

In our studies, the agreement between the enriched GO terms to the corresponding protein lists has not always been the same. From our past study of housekeeping genes ^40^, a disagreement has stood out. In that study, we have discovered that in 13 published housekeeping gene lists by pan-human organ/tissue transcriptomics analyses, there was only a single gene in common. Yet the corresponding enriched GO biological processes were surprisingly uniform and consistent. We had advocated “housekeeping functions” over “housekeeping genes” ^40^ due to the unanimity in the enriched biological functions and the disparity in the identified housekeeping genes. Even though in sharp contrast of the GO terms derived between the housekeeping genes and the OSPs as we shown here, both results suggest that Gene Ontology can well reflect the overall state of the identified proteins and can be used as a reliable criterion to evaluate the nature of the obtained transcripts and proteins.

## CONCLUSIONS

We presented here a new experimental scheme to study organ specific proteins (OSPs) that we named it breadth study. In this scheme, we expanded the coverage of diverse genetic backgrounds balanced by decreased analysis depth as opposed to the existing in-depth studies. Our results revealed large diversity in OSPs in mice of different genetic backgrounds, and we used the robustness score (R) to catalog OSPs that are consistent across genetic backgrounds from those are varied. Depending on the sampling depth, the number of OSPs and their derivation process, their robustness can vary substantially. Nevertheless, at all abundance levels filtering OSPs for consensus among genetic backgrounds by the breadth study can effectively purify OSPs. Furthermore, we discovered that biological functions enriched by the corresponding OSPs are also organ specific, and sensitive to the proteins from which they are derived. We defined D score here to quantitatively evaluate the organ specificity at both protein level and function level. A higher D score indicates a higher organ specificity and less overlap to other organs.

Finally, linking our observed diversity in mouse strains and those made in gene expression analyses of eQTL and pQTL ^13-15^, to the clinical knowledge of organ specific biomarkers in patients ^19,41^, we emphasize the importance to address such dynamic changes of OSPs in a population. The analysis here demonstrated that a single nanoLC-MS/MS run carried out by modern high accuracy and high sensitivity tandem mass spectrometry within 1-2 hours can provide sufficient protein identifications that enable fast and practical breadth studies. It allows the derivation of a relatively large number of OSPs to quantitatively distinguish them into relatively small subcategories based on their R and D scores for various clinical as well as biomedical applications. Comparing with the existing depth studies, the breadth analysis extends an orthogonal dimension with comparable resources, simplified statistics, and a shortened distance to clinical utility. More importantly, such study can be easily scaled to include more strains and conditions due to its nature of label free. We hope the strategy and the lists of the filtered OSPs can benefit the medical and biological research in organ specific diagnoses and treatments.

## ACKNOWLEDGEMENTS

This work is supported by the Canada Foundation of Innovation, the British Columbia Knowledge Development Fund, the Stem Cell Network in Canada, Simon Fraser University, the United States Department of Defense grants of W911SR-06-C-0057 and W911SR-07-C-0101, and the National Sciences and Engineering Research Council of Canada. Y.Z. would like to acknowledge the financial support from China Scholarship Council and Mitacs Globalink program.

## AUTHOR CONTRIBUTIONS

L.H. B.S. Z.H. conceived the idea, C.L. S.Q. G.L. A.U. P.F. B.S. Z.H. performed the experiments, B.S. Z.S. A.F. Z.Y. K.L. analyzed the data, K.W. R.M. contributed to the project administration, B.S. and L.H. wrote and everyone reviewed the manuscript.

