## Supplementary text for "Mouse Organ Specific Proteins and Functions"

### TABLE OF CONTENTS

**Methods**

1. Tissue histology

2. Proteomic sample processing

3. MS analysis of peptides.

4. MS/MS database search parameters and protein inference

**Supplementary Tables**

**Table S1:** Total detected proteins.

**Table S2:** Verified Organ Specific Proteins (vOSPs).

**Table S3:** Top10 enriched biological processes of Verified Organ Specific Proteins.

**Table S4:** Complete Organ Specific Proteins (cOSPs)

**Table S5:** Top10 enriched biological processes of Complete Organ Specific Proteins.

**Table S6:** Conditional OSPs from top100 organ proteomes.

**Table S7:** Top10 enriched biological processes of conditional OSPs.

**Methods:**

*Tissue histology.* Sections of the excised organs were stored in 10% formalin for H&E staining in the Department of Pathology, University of Washington, Seattle, WA. General steps included the embedding of tissues in paraffin, the slicing and mounting on the microscope slides, staining with Hematoxylin and Eosin (H&E staining), prior to microscopic imaging.

*Proteomic sample processing.* Frozen organs were pulverized in ceramic crucible chilled with liquid nitrogen and collected to Eppendorf tubes. PBS with 1:100 diluted protease inhibitor cocktail (PIC) was used to rinse the tissue on ice until the majority of blood had been removed. Then samples were transferred to PBS with 1:100 PIC and 2-5% SDS, before homogenized by Precellys24 (Bertin, France) at 4 °C. The supernatant was obtained by centrifugation and the protein contents were quantified by BCA. Fifty microliter of samples were processed for digestion. In detail, TCEP and EDTA were introduced to the final concentration of 10-mM and 5 mM respectively, and samples were boiled at 100°C for 10 min. Afterwards, samples were processed based on the filter aided sample preparation (FASP) method [1], in which 8M Urea was introduced and SDS was removed by filter through centrifugal force. The denatured and reduced proteins were alkylated by sodium iodoacetamide and quenched by DTT, before introducing sequence grade trypsin at 1:50 enzyme to protein ratio and digesting at 37 °C overnight. The digested sample were desalted on a Sep-Pak C18 column and dried in a SpeedVac® (Thermo Savant, Holbrook, NY, USA) concentrator.

*MS analysis of peptides.* Three different systems were used, i.e. LTQ XL linear ion trap mass spectrometer (LTQ), Velos Pro dual-pressure linear ion trap mass spectrometer (Velos), and LTQ-Orbitrap XL hybrid ion trap-Orbitrap mass spectrometer (Orbi). For LTQ analysis, similar to previous published procedure [2,3], an in-house fabricated nanoelectrospray source and an HP1100 solvent delivery system (Agilent Technologies) were coupled to the mass spectrometer. Samples were automatically delivered by a FAMOS autosampler (LC Packings, San Francisco, CA) to a 100-μm (ID) fused silica capillary precolumn packed with 2 cm of 200-Å pore size Magic C18AQTM material (Michrom Bioresources, Auburn, CA). The samples were washed with solvent A (5% acetonitrile in 0.1% formic acid) on the precolumn, eluted with a gradient of 10–35% solvent B (100% acetonitrile) over 60 min to a 75-μm and 10-cm fused silica capillary column packed with 100-Å pore size Magic C18AQ material (Michrom Bioresources), and then injected into the MS at a constant flow rate of 300 nL/min. Eluting peptides were analyzed by tandem MS in data-dependent acquisition, in which five most abundant precursor ions were selected for MS2 fragmentation with a dynamic exclusion of 1 repeat count in 30 sec for 180-sec exclusion, and the size of exclusion list is 50. For Orbi analysis, an Agilent 1200 HPLC was coupled to the instrument. Peptides were applied to a fused and fritted silica capillary pre-column of 100 μm ID (New Objectives, Woburn, MA) packed with 2 cm of 200 Å pore-size C18 resin. Samples were subsequently washed with solvent A (5 % acetonitrile in 0.1 % formic acid) on the pre-column, eluted with a gradient of 10-35% solvent B (100% acetonitrile) over 60-90 minutes to a 75 μm x 10 cm fused silica capillary column packed with 100 Å pore-size Magic C18AQ™ (Michrom Bioresources, Auburn, CA), and then injected into the Orbi with a nano-ESI source. Eluting peptides were analyzed in MS by similar data-dependent acquisition as before with a dynamic exclusion setting of 1 repeat count in 30 sec for 60-sec exclusion, and the size of exclusion list is 50 (60-min gradient) or 500 (90-min gradient). Majority of samples were analyzed by Orbi method. For Velos analysis, Eksigent nano-HPLC was used with similar columns and 60-min gradient LC method as Orbi, and similar data-dependent acquisition was carried out by Velos with minor modifications, in which top 10 most abundant precursor ions were selected for MS/MS fragmentation with a dynamic exclusion setting of 1 repeat count in 30 sec for 180-sec exclusion, and the size of exclusion list is 50.

*MS/MS database search parameters and protein inference.* For Orbi and Velos runs the parent mass error was set to ± 50 ppm and for LTQ runs, it was set to ±3.0 Da. For MS2, 0.5 amu was used for all analyses. For all runs, peptides were allowed to be semitryptic with up to two internal cleavage sites. The search parameters included a fixed modification of +57.0215 for carbamidomethylated cysteines and a variable modification of +15.9949 for oxidized methionines and +42.0106 for protein n-terminal acetylation.
